# Plasmidome-defensome interactions drive adaptation of ‘*Fervidacidithiobacillus caldus’* in natural and industrial extreme acidic environments

**DOI:** 10.1101/2025.04.02.646953

**Authors:** Sebastián Pacheco-Acosta, Gustavo Castro-Toro, Camila Rojas-Villalobos, Cesar Valenzuela, Juan José Haristoy, Abraham Zapata-Araya, Ana Moya-Beltrán, Pedro Sepúlveda-Rebolledo, Ernesto Pérez-Rueda, Ricardo Ulloa, Alejandra Giaveno, Francisco Issotta, Beatriz Diéz, Simón Beard, Raquel Quatrini

**Affiliations:** Facultad de Medicina y Ciencia, Universidad San Sebastián, Santiago, Chile; Centro Científico y Tecnológico de Excelencia Ciencia & Vida, Santiago, Chile; Facultad de Ingeniería, Arquitectura y Diseño, Universidad San Sebastián, Santiago, Chile; Departamento de Informática y Computación, Facultad de Ingeniería, Universidad Tecnológica Metropolitana, Santiago, Chile; Instituto de Investigaciones en Matemáticas Aplicadas y en Sistemas, Universidad Nacional Autónoma de México, Unidad Académica del Estado de Yucatán, Carretera Sierra Papacal, Mérida 97302, Yucatán, México; PROBIEN (CCT Patagonia Confluencia-CONICET, UNCo), Facultad de Ingeniería, Departamento de Química, Universidad Nacional del Comahue, Neuquén, Argentina; Centro GEMA - Genómica, Ecología & Medio Ambiente, Universidad Mayor, Chile; Millennium Institute Center for Genome Regulation (CGR), Santiago, Chile; Center for Climate and Resilience Research (CR)2, Chile; Departamento Genética Molecular y Microbiología, Facultad de Ciencias Biológicas, Pontificia Universidad Católica, Santiago, Chile

**Author notes:** **Correspondence:** Raquel Quatrini, Simón Beard. Join First Authors (Sebastián Pacheco-Acosta, Gustavo Castro-Toro).

**Keywords:** *Acidithiobacillia*, *Acidithiobacillus*, mobile genetic element, MGE, plasmid, defense system, Restriction-Modification system, CRISPR-Cas system, plasmid ecology

## Abstract

Plasmids are major drivers of microbial evolution, enabling horizontal gene transfer (HGT) and facilitating adaptation through the dissemination of relevant functional genes and traits. However, little is known about plasmid diversity and function in extremophiles. ‘*Fervidacidithiobacillus caldus’,* a meso-thermo-acidophilic sulfur oxidizer, is a key player in sulfur cycling in natural and industrially engineered acidic environments. Here, we present a comprehensive analysis of the plasmidome, and associated anti-mobile genetic element (anti-MGE) defense systems (defensome), across genomes of this species and metagenomes from diverse natural and industrial settings harboring ‘*F. caldus’*. We identified >30 distinct plasmids, representing five consistent replication-mobilization families. Plasmids ranged in size between 2.5–65 kb, with gene content and plasmid modularity scaling with element size and copy numbers inversely correlating with size. Plasmids carried variable numbers of hypothetical proteins and transposases, with annotated cargo genes reflecting functional differentiation by habitat. Defensome profiling revealed over 50 anti-MGE systems in sequenced ‘*F. caldus’* isolates, including diverse restriction-modification systems, CRISPR-Cas types IV-A and V-F, and widespread abortive infection and composite defense systems such as Wadjet, Gabija, and Zorya. In environmental populations, an inverse relationship was observed between defensome complexity and plasmidome abundance and diversity, underscoring a pivotal role of the host defensome in modulating persistence, compatibility, and overall plasmid diversity across ‘*F. caldus’* populations. Yet, other plasmids appeared decoupled from both host abundance and defensome complexity, suggesting potential host shifts, environmental persistence, or differential replication under suboptimal growth conditions for the host. Altogether, these findings reveal a modular, adaptive plasmidome shaped by selective pressures and host–plasmid–defensome interactions and positions plasmids as key contributors to adaptation, gene flow, and functional innovation in this extreme acidophile.

**Importance:** Plasmids are key vehicles of gene exchange and adaptation in bacteria, yet their roles in extremophilic systems remain poorly understood. This study provides the first integrated view of the plasmidome and defense systems in ‘*Fervidacidithiobacillus caldus’*, a sulfur-oxidizing acidophile relevant to both natural biogeochemical cycling and industrial bioleaching. We uncover a rich plasmid diversity structured into modular families with variable cargo and backbone features and reveal their coexistence with complex anti-MGE defense repertoires. By combining genomic and metagenomic approaches, we expose principles of plasmid compatibility, persistence, and habitat-specific adaptation. These insights expand current knowledge of mobile genetic elements in extreme environments and provide a foundation for plasmid-based vector design and synthetic biology in acidophiles, with direct implications for biomining and environmental remediation in extreme environments.

## Introduction

Plasmids are pivotal elements in bacterial evolution and adaptation, particularly through horizontal gene transfer (HGT), which enables the exchange of genetic material between distantly related bacteria within microbial communities [1,2]. HGT mediated by plasmids is a major driver of bacterial diversification, enabling the rapid acquisition of adaptive traits, such as antibiotic resistance, which spread rapidly under anthropogenic pressures [3]. Plasmids also promote adaptation by increasing gene dosage, enhancing the potential for mutation and diversification of their cargo, and giving rise to phenomena such as heteroplasmy [4], heterozygosity [5], and genetic dominance [6]. These processes can guide the expansion of populations under changing environmental conditions and accelerate bacterial evolution. The plasmid repertoire of prokaryotes is vast and highly diverse, both in terms of adaptive cargo and core backbone genes (e.g. replication and mobilization modules), which vary across hosts and environments [7]. Substantial variability has been documented in plasmid genome size, architecture, and topology [8], as well as in copy number and host range, mirroring the diversity of their prokaryotic hosts. Plasmids are ubiquitous in nature, being found in both conventional environments such as soil and water, and in extreme habitats such as geothermal sites and hypersaline ecosystems [9,10]. However, despite their ubiquity, most plasmid research has centered on model microorganisms of medical or industrial relevance, leaving plasmids in extremophiles comparatively understudied. This gap stems in part from the challenges of cultivating and genetically manipulating extremophiles, whose specialized growth requirements and slow replication rates limit experimental tractability and hinder large-scale functional analyses. Yet, studying plasmids in these organisms remains critical to uncover unique adaptive traits and molecular strategies underlying extremophile adaptation and resilience, while also providing genetic parts and design principles valuable for synthetic biology.

Among extremophiles, members of the class *Acidithiobacillia* are obligate chemolithoautotrophs and extreme acidophiles that thrive in highly acidic, sulfur- and metal-rich environments. ‘*Fervidacidithiobacillus caldus’* (formerly *Acidithiobacillus caldus*), a well-studied member of this class, grows optimally at 40-45°C and pH ∼2.5 [11]. Strains of this species have been isolated from diverse geographical locations and habitats [12], reflecting its broad adaptability to various abiotic and biotic conditions. Genomic surveys of ‘*F. caldus’* have revealed moderate genetic diversity among isolates [12–14], largely attributed to horizontal gene transfer of mobile genetic elements, including plasmids [15–17]. Yet, the adaptive value of plasmid-encoded functions has not been tested experimentally, and little is known about their distribution, maintenance, or potential fitness costs at the populational level [18]. Such costs could be particularly detrimental for extremophiles, which typically display starvation-survival strategies, including slow growth, low cell densities, reduced biomass yields, and high degrees of spatial isolation [19].

In addition to this, an increasing suite of defense systems acting on plasmids has been described in recent years in different model microorganisms [20–24], including systems that inhibit plasmid replication or transmission, and limit HGT of their cargo genes [25,26]. Thereafter, the persistence of plasmids in host cell populations is shaped by a series of interacting factors that sustain the plasmid life cycle [27], including plasmid-encoded traits (e.g. replicon type), host factors (e.g. Restriction-Modification systems [28]), and incompatibilities with co-occurring MGEs [29], all further influenced by environmental selection acting on the accessory genes carried by the plasmid.

Despite renewed interest in plasmid ecology and evolution (fueled by the expansion of metagenomic datasets) their contribution to the structuring of microbial communities and to HGT *in situ* is still poorly understood, particularly in acidic environments. This study investigated the repertoire of plasmids (plasmidome) and defense systems (defensome) of ‘*F. caldus’* sequenced strains and populations in both natural and industrial acidic environments. By analyzing the occurrence, diversity, and distribution of plasmids, the nature of their backbone gene modules and cargo genes, and associated defense systems, we aimed at elucidating how plasmids contribute to the adaptation of ‘*F. caldus’* to its environment, and how defense systems influence plasmid persistence and gene flow. Understanding these dynamics is crucial for advancing knowledge of extremophilic biology and could inform biotechnological applications involving bioleaching and bioremediation processes.

## Results and discussion

Novel plasmids and variants of the 8 known plasmids were successfully identified in 13 out of 17 strains of the species sampled globally (**Fig.1a, Table.S1a**), with a general plasmid prevalence of >75% (**Fig.1a)**. A similarly high prevalence of plasmids (in the range of 70– 85%) has been reported in other bacteria (e.g.[30]), yet according to current understanding is higher than in other *Acidithiobacillia* class species [31]. The identified replicons varied in size, ranging from 2.6 to 65 Kb, with a median size of 14 Kb (**Fig.2a**) and had a relative abundance with respect to the cognate chromosomal replicons of 0.6 to 6.2 fold (**Table.S2**). The calculated plasmids copy numbers correlated negatively with the size of the elements (**Fig.2b**). This suggests that the candidate secondary replicons of the *Acidithiobacillia* would all be plasmids with a medium to low copy number, as is indeed the case for experimentally evaluated plasmids of the class [32]. With few exceptions, the % G+C content of all these plasmids was lower than that of the respective host genomes (**Table.S2**).

**Figure 1.**
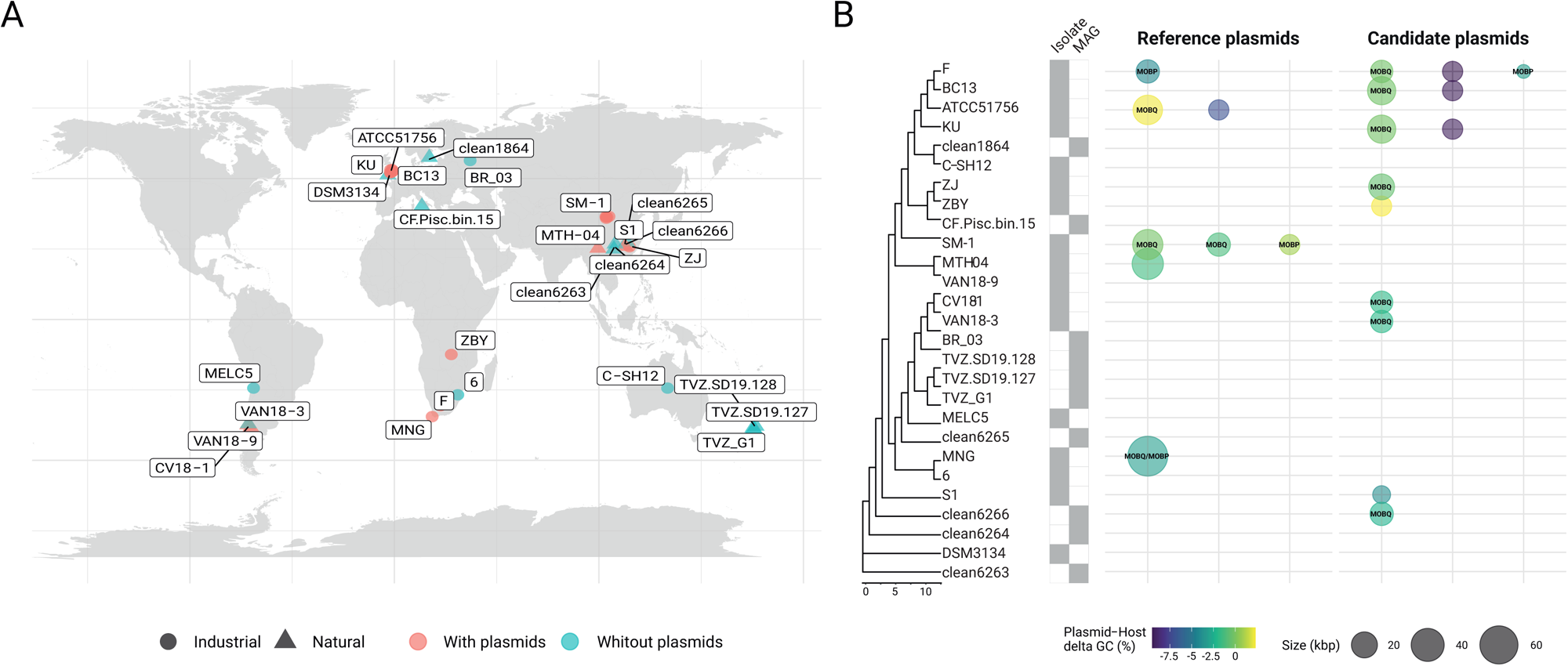
Prevalence and phylogenetic distribution of plasmids in ‘*F. caldus’* genomes. (**A**) Geographical origin and source type (natural or industrial) of ‘*F. caldus’* isolates and MAGs included in this study. Strains harboring plasmids are shown in light red, while plasmid-free strains are shown in light blue. (**B**) Maximum-likelihood (ML) phylogenetic tree based on the concatenated alignment of 120 bacterial marker proteins (*bac120*; [94]) from 17 isolate genomes and 10 MAGs available in GenBank as of July 2024. Alignments were generated with MUSCLE, and ML-tree inference was performed using IQ-TREE v2.0 with the JTT substitution model, a discrete Gamma distribution, and 100 bootstrap replicates, and the MEGA X suite v10.2.3 [116]. *Thermithiobacillus tepidarius* DSM3134 T was used as outgroup to root the tree. Plasmid occurrence is represented by a dot: dot size reflects known or inferred plasmid size, and color indicates the difference in G+C content between the plasmid and the host chromosome. For mobilizable plasmids, the type of relaxase (as classified by [107]) is indicated inside the dots.

**Figure 2.**
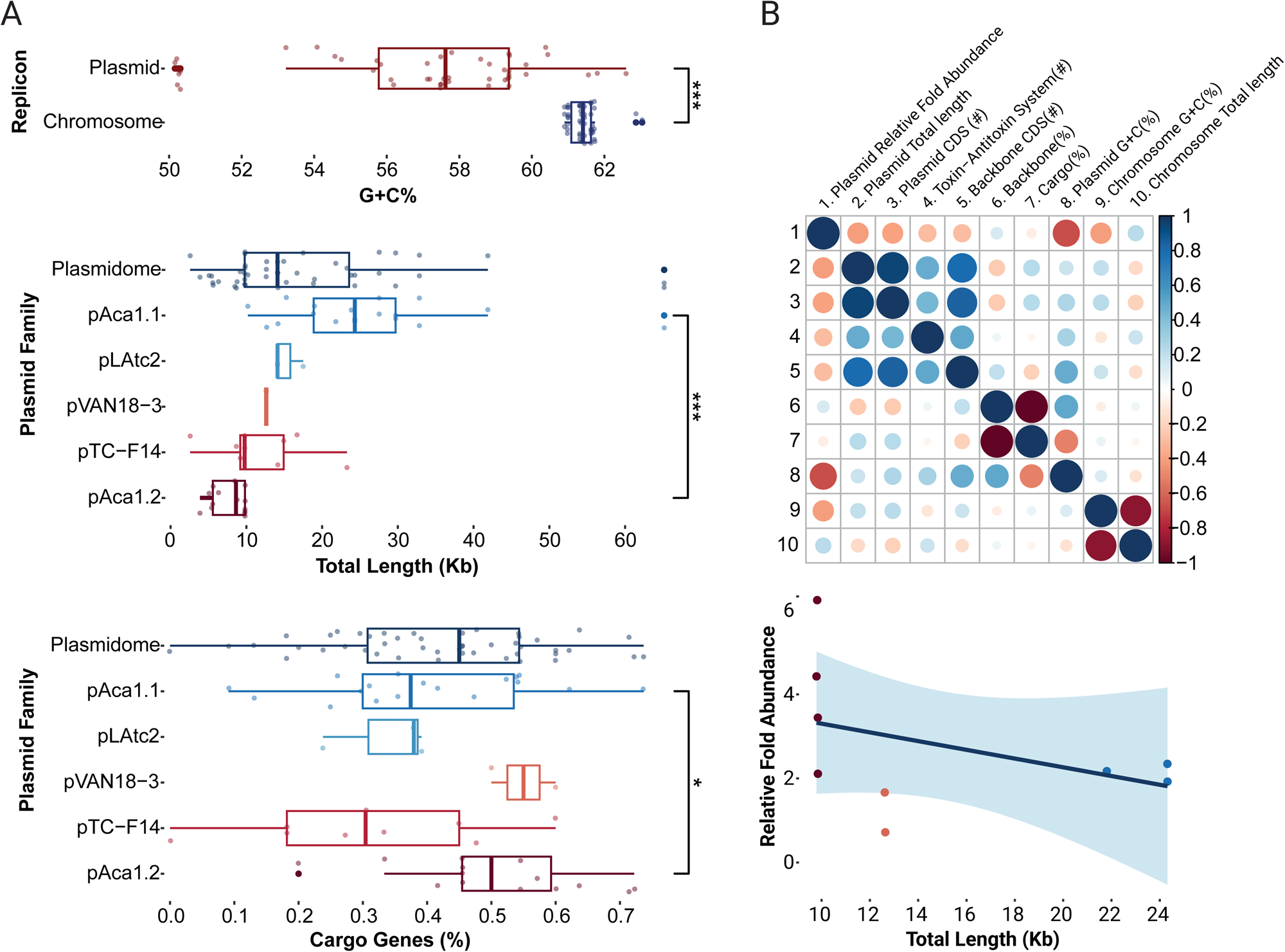
Distribution of genomic features across ‘*F. caldus’* plasmids. (**A**) Boxplot of the distribution of plasmid G+C content, size (kb), and proportion of cargo genes. Plasmids GC content (%) is significantly lower G+C than their cognate chromosomes (***P* < 0.001**) and vary widely in size (2.6–65 kb; median: 14 kb). Significant differences were also observed in plasmid size and cargo gene content across plasmid families (*P < 0.001*\* and *P* < 0.05, respectively). (**B**) Correlation analyses between plasmid genomic metrics. The correlogram (top) shows Pearson correlation coefficients between selected features, with blue indicating positive correlations and red negative. Color intensity reflects correlation strength. The scatterplot (bottom) illustrates the inverse relationship between plasmid size and inferred plasmid copy number (trend line and confidence interval), based on normalized read depth with respect to the chromosome, and indicated here as relative fold abundance.

The *in silico* analysis of publicly available ‘*F. caldus’* genomes (**Table.S1a**) revealed that the number of plasmids varied from one to three plasmids per strain, with 35.7% of the strains harboring one plasmid, 71.4% multiple plasmids, and 17.6% being devoid of plasmids. Even if lower than the plasmid carriage (8-10 plasmids per strain) observed in clinical and environmental bacteria [33–35], including certain acidophiles (e.g., 8 plasmids in *Acidiphilium* spp. [36]), the co-carriage of multiple plasmids per isolate is of significance particularly in the light of the limited evidence for plasmid incompatibility in ‘*F. caldus’* [37] or in acidophiles in general [31].

To determine the basis of their coexistence and obtain hints on the fitness costs of the plasmidome, we classified ‘*F. caldus’* plasmids into distinct families based on their core gene content (**Table 1**). Given that plasmid incompatibility (defined as the inability of two co-resident plasmids to be stably inherited by daughter cells in the absence of selection relies on features of the replicon, such as the origin of replication and/or the partitioning system [38]), we first analyzed these aspects across the recovered plasmids. Sequence-based clustering of plasmid-encoded proteins revealed 351clusters, with 40% corresponding to conserved backbone protein families (PFs) and 60% representing accessory protein families, several of which conformed distinct gene modules (**Fig.3**). Sequence analysis identified five distinct replication modules, differing in replicase type (single or multidomain, **Fig.3a**) and replicon organization (**Fig.3b**). All modules harbored a variant of the replicative helicase (**Table.S2**). The *rep* gene from plasmids represented by pTC-F14 (RepA_1, 291 aa) and pVAN18-3.1 (RepA_2, 435 aa) encoded distinct proteins with sequence similarity to the replicative helicase of the RSF1010 plasmid family (pfam13481, [39]). The pTC-F14 replicase (RepA_<u>1</u>; IncQ-2b) shares with other IncQ-family replicons the presence of three replication related genes, *repB, repA* and *repC*, encoding the primase, the helicase and the iteron-binding protein, respectively [40,41], in the proximity of a 22-bp iteron based *ori*V [32]. pTC-F14-like plasmids were detected in strains MNG, SM-1 and S1 (pTcM1.1, pLAtc1, pS1, pCanu) with conserved organization and *ori*V, supporting a rolling-circle replication mechanism [42] with medium copy numbers (12–16 copies per chromosome [32]). In contrast, pVAN18-3.1 (RepA_2) lacked homologous primase and iteron-binding proteins but encoded a large multidomain replicase (> 400 aa) and a small (132-135 aa) DNA binding protein, suggesting functional divergence. The latter showed similarity to ORF2 (NF038291) from the replication region of plasmid pAB02 and related *Acinetobacter* plasmids, supporting a functionally divergent replication strategy for these medium sized plasmids. The replicase from pAca1.1-like (RepA_3, 437 aa) and pAca1.2-like (RepA_4, 335 aa) plasmids encoded a three-domain protein containing a replicase, a primase and a C-terminal DNA-binding domain and were found adjacent to a predicted *ori*V and to a replication associated CDS encoding either KfrA or an ortholog of ORF2 (pAB02_ORF2). The fifth type of replication module uncovered in ‘*F. caldus’* plasmid pLAtc2 (RepA_5, 329 aa) encoded a RepA_C (pfam04796) that is only 28.6% similar to its best matching hit in ‘*F. caldus’* (RepA, pTC-F14). This replicase was encoded next to the *ori*V and convergent to a KfrA-encoding gene, a plasmid-specific nucleic acids binding protein (NAP) found also in other plasmids of the *Acidithiobacillia* class [43], and recently shown, in the conjugative plasmid RA3 of *Pseudomonas aeruginosa*, to mediate low-copy-number plasmid segregation during cell division through interactions with the segregosome proteins IncC (ParA) and KorB (ParB [44]. These rep-modules distributed differentially among strains of the species (**Table.S3**), with several evident plasmids concurrencies or exclusions, underscoring compatibility between replicons Rep1:Rep3, Rep1:Rep5 and Rep3:Rep4 and incompatibility between Rep3:Rep5 and Rep2:all (**Fig.S1a**). Based on the occurrence of a single replicon-type per strain, all these replicons are inferred to be self-incompatible, except for Rep4 found in variant plasmids related to pAca1.2 in several strains (**Fig.S1b**). Also, one plasmid carrying two replicons could be identified as a cointegrate of a pTC-F14-like plasmid (RepA_1) and a pAca1.1-like (RepA_3), with evidence of compromised integrity of the *repABC* replicon (pTcM1 in strain MNG; [45]).

**Figure 3.**
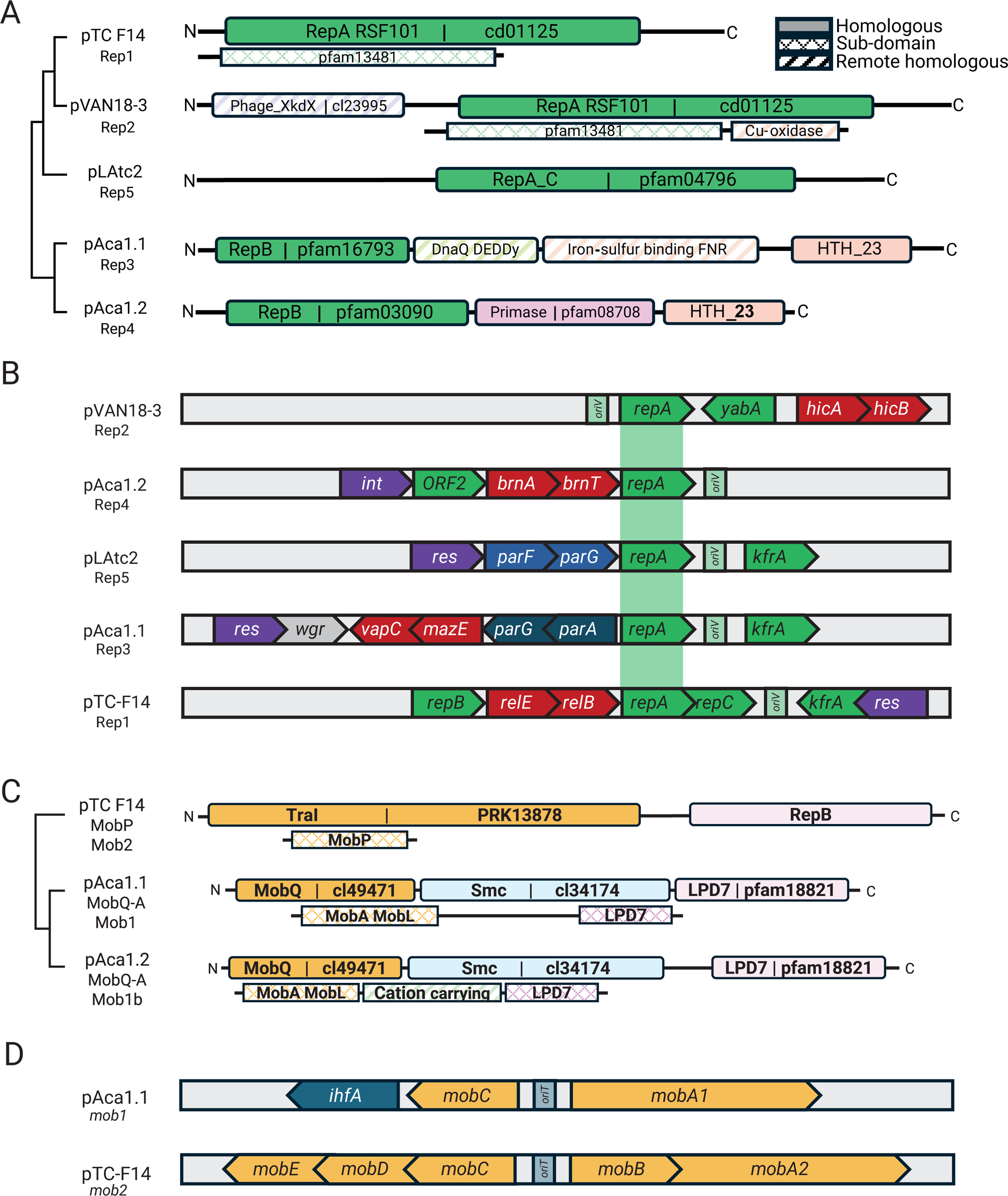
Backbone module organization of ‘*F. caldus’* plasmids. (**A**) Replication modules RepA_1 to RepA_5 identified across ‘*F. caldus’* plasmids, differing in replicase type (single- or multi-domain) and associated protein domain architecture. (**B**) Gene vicinity of RepA per replicon type, showing adjacency of replication (green), stabilization (red) and partition genes (blue) and the replication origin location (*ori*V region). (**C**) Clustering by mobilization module types (*mob1* vs. *mob2*) and protein domain architecture of identified relaxases (MOBQ and MOBP_A families) in ‘*F. caldus’* plasmids. (**D**) Gene architecture of mobilization modules *mob1* and *mob2*, highlighting differences in relaxase family, accessory proteins (RAPs), and *ori*T region location. Gene architectures support the classification of plasmids into functional families and offer insights into compatibility and inheritance mechanisms.

**Table 1.** Plasmid families identified in ‘*F. caldus’* sequenced strains and MAGss.

| Plasmid family | Representative plasmid | Size range (Kb) | Relaxase MOB type | 1ry RepA type | 2ry RepA type | PAR system | TA systems (#) | Total genes | Cargo genes (#) | Cargo genes w/ function (#) | Backbone (%) | Concurrence per strain |
| --- | --- | --- | --- | --- | --- | --- | --- | --- | --- | --- | --- | --- |
| 1<br>(n=5) | pTC-F14 | 9.8-14,2 | PA | Rep1 | - | - | 1 | 21 | 10 | 4 | 52,4 | 3, 4, 5 |
| 1, 3<br>(n=1) | pMNG_TcM1 | 65,2 | Qa, PA | Rep3 | Rep1 | Par1 | 3 | 103 | 44 | 23 | 35,9 | 1,3 |
| 3 (n=13) | pAca1.1 | 27.5 - 32.8 | Qa | Rep3 | - | Par1 | 2 | 39 | 21 | 11 | 46,2 | 1, 4, 5 |
| 5<br>(n=2) | pLAtc2 | 14.1 | Qa | Rep5 | - | Par2 | 0 | 20 | 5 | 3 | 75,0 | 1, 3, 4 |
| 2<br>(n=2) | pVAN183_JAAOM N010000091 | 12.6 | Qa | Rep2 | - | - | 1 | 13 | 7 | 4 | 46,2 | - |
| 4a<br>(n=5) | pAca1.2 | 9.8 | - | Rep4a | - | - | 1 | 11 | 6 | 3 | 45,5 | 1, 3, 4, 5 |
| 4b<br>(n=4)* | pS1_LZH01000935 | 5.6-8.5 | - | Rep4b | - | - | 1 | 11 | 7 | 2 | 36,4 | 1, 3, 4a, 4c, 4d, 5 |
| 4c<br>(n=2) | pZJ_LZYG01000035 | 5.1-8.8 | - | Rep4c | - | - | 2 | 12 | 5 | 3 | 58,3 | 3, 4b, 4d |
| 4d (n=2) | pZBY_LZYF01000110 | 3.9-5.4 | - | Rep4d | - | - | 1 | 7 | 4 | 3 | 42,9 | 3, 4b, 4c |

**Abbreviations:** n: number of strains/genomes/MAGs carrying the plasmid; (\*): integrated in a chromosomal contig.

Partitioning systems were consistently located adjacent to replication genes, forming seamlessly integrated modules (**Fig.3b**). Sequence analysis identified ParFG-*parH*-like systems in ‘*F. caldus’* plasmids [46], resembling those of the multiresistance plasmid TP288 [47], and comprising four ParF (COG1192; pfam13614) and three ParG variants. ParF proteins shared 37.7–83.8% identity among ‘*F. caldus’* plasmids and 41.9–52.6% with ParF from TP288, while ParG orthologs were more divergent (23.5–32.2%), suggesting adaptive differentiation of centromere-binding factors. A conserved 4-nt tandem repeat (5’-CTAT-3’), flanked by AT-rich sequences, was also identified near the *parFG* loci and likely represents a centromere-like site in pAca1.1-like plasmids. The Par system components were consistently associated with the pAca1.1 plasmid family, the most widespread in ‘*F. caldus’* (12 strains and 2 MAGs), supporting the notion that partition is the most important determinant of plasmid stability [47]. However, their presence alone does not guarantee compatibility, as co-occurrence of plasmids with different Par variants was observed in only one case (strain F: pAca1.1-like [JAAOML010000407] and pTC-F14 [JAAOML010000340]). Since partition systems are ubiquitous in low-copy number plasmids (or generally absent in high-copy number plasmids), we infer that all pAca1.1_like plasmids are low copy number plasmids. This aspect is supported by plasmid copy number inference (1.67 plasmids/per chromosome; **Table.S2**) and experimentally proven to be low (1 copy of pAca1.1 every 2 chromosomal copies) in the ‘*F. caldus’* ATCC 51756^T^ [48].

To improve the classification of plasmid families in ‘*F. caldus’*, we characterized their gene modules responsible for dispersal and stabilization within hosts. A total of 13 unique relaxases were identified, belonging to MOBQ or MOBP_A families [49], with MOBQ relaxases being the most ubiquitous in the dataset (**Fig.3c**). Relaxases clustered into two mobilization module types, *mob1* and *mob2* (**Fig.3d**). The *mob1* modules (8 variants), encoded a MobA relaxase of the MOBQa type, along with three small relaxase-accessory proteins (RAPs). These RAPs include a MobC ortholog, an IHF histone-like protein, and a DNA-binding protein (HTH_23:PF13384; DUF742: PF05331) of unclear function. Presence of IHF coding genes and IHF-binding sites in the *ori*V region of *Acidithiobacillia* class plasmids has been previously reported [43, 50–51] and inferred to have a role in DNA-bending linked to plasmid replication and/or transfer facilitation [52, 53]. In turn, *mob2* modules (5 variants) encoded a MobA relaxase of the MOBP_A type and four conserved accessory proteins (MobBCDE). The *mob1* module was found in all pAca1.1-related plasmids (n=11, Rep3-type) and pLAtc2-like plasmids (n=2, Rep5-type), which are all medium to large plasmids, while the *mob2* module was invariably linked to smaller-sized IncQ2 plasmids (n=5, Rep1-type), including well-characterized pTC-F14 [54]. In addition to the relaxase domain (pfam03432) of the MOBP_A family, the MobA_mob2 proteins contained a second conserved domain present in RepB primases (pfam16793) and clustered with 4 defined, yet variable, Mob accessory proteins (sequence-level similarity range: 21.2-77.8%). Despite this variability, their conserved occurrence and organization reinforce previous findings on the essential role of *mobA, mobB*, and *mobC* RAPs in pTC-F14 plasmid mobilization in ‘*F. caldus’* [54]. The MobA-RepB fusion proteins displayed a high degree of conservation, with 71.3%–100% sequence similarity among ‘*F. caldus’* plasmids and 73.0%–89.2% similarity to its only known homolog in the *Acidithiobacillia* class, the pTF-FC2 plasmid. Probable origins of transfer *ori*T could be predicted in all plasmids, at the intergenic region between the *mobA* gene and divergently transcribed RAPs (**Fig.S2**). The sequence of the *ori*T was highly conserved between plasmids with the same *mob*-type (%I), yet not between plasmids pertaining to different mob types (%I). Still all *ori*Ts conformed to a general palindromic configuration described previously for plasmids pTF1 and pTC-F14, being adjacent to a highly conserved predicted nick site. All ‘*F. caldus’* plasmids lacked conjugation genes, yet the widespread occurrence of mobilization genes indicate they necessarily co-opt the type IV secretion machinery of other self-transmissible MGEs [55–56].

Twenty nine out of thirty plasmids in the ‘*F. caldus’* plasmidome encoded at least one toxin-antitoxin (TA) system, with 46.7% carrying multiple systems (**Table.S2**). These were invariantly type II TA systems, which are frequently associated with MGEs and HGT in other organisms [57]. Five toxin families were identified: RelE, VapC, HicB, BrnT, and Doc, each with 1–4 variants. RelE protein family was the most widespread, with multiple subtypes (4 RelE, 1 HigB, 1 YafQ). In agreement with general knowledge [58], TA operons were typically bicistronic, consisting of toxin-antitoxin gene pairs, (e.g., *relBE, brnTA*, etc.); an exception was pLAtc1, which harboured non-cognate *relE* and *vbhA* genes in separate loci. TA systems were frequently located between plasmid backbone genes related to replication (*repA, kfrA*), mobilization (*mobA, mobE*), or partitioning (*parG*), reinforcing their role in plasmid stabilization. Two organizational patterns were observed: canonical (*antitoxin-toxin*, e.g., *vapBC, relBE, dinJ-yagQ, hicBA, phd-doc*) and non-canonical (*toxin-antitoxin*, e.g., *brnT-brnA, higB-higA*) [59]. Interestingly, the non-canonical configuration was found exclusively in smaller plasmids (e.g., pAca1.2-like plasmids carrying *brnT-brnA*), possibly reflecting an aspect of the regulation of compact replicons. However, specific studies investigating the effects of reversed gene orientation on the maintenance efficiency of small versus larger plasmids are unavailable. All TA toxins in ‘*F. caldus’* targeted translational processes [60]; they are predicted to function as ribonucleases (*RelE* targets mRNA, *HicA* degrades mRNA independently, *VapC* cleaves initiator *tRNAfMet*), translation elongation inhibitors (*Doc* phosphorylates EF-Tu, disrupting translation), or act on undefined RNA targets (*BrnT*). Larger plasmids frequently combined 2 or 3 TA systems with different specificities (*pAca1.1*: *vapBC* [tRNA] and *relBE* [mRNA]). Also, plasmids coexisting in the same strain tended to encode distinct TA types (pAca1.1 [vapBC, relBE] *vs.* pAca1.2 [brnTA]), or variants within the same type (pF_JAAOML010000123 pAca1.1-like [relE1] *vs.* pF_JAAOML010000310 pTC-F14-like [relE2]), suggesting independent plasmid stability control. This strategy likely enhances plasmid persistence while preventing exclusion of competing plasmids with similar backbone structures [61].

Insights into the organizational principles of the ‘*F. caldus’* plasmidome were obtained from the plasmid metadata and annotations (**Table 1**). Plasmids segregated into two main size classes with distinct genetic features. Small plasmids (<10 kb), exclusively harbored Rep4-type replicons and non-canonical TA systems, lacked mobilization and partition modules, and were mostly cryptic, with few or no functionally annotated genes. Despite their minimal gene content, these plasmids were detected in >50% of strains and variant types often coexisted in certain strains with up to three variants in some cases (e.g. ZJ/ZBY). Some appeared integrated into the chromosome or larger MGEs (e.g., var4b), consistent with the presence of FimB-FimE-Int type integrases in some pAca1.2-like plasmids. Coverage analysis further indicated these small plasmids were present at significantly higher copy numbers (3-6x chromosome levels) than coexisting larger plasmids, suggesting dosage-dependent roles. Although functionally uncharacterized, their persistence and widespread distribution across strains indicates they are probably more than neutral genomic passengers. In contrast, ‘*F. caldus’* larger plasmids (>10 kb) exhibited greater functional complexity, incorporating partitioning, mobilization, and stabilization modules, supporting long-term maintenance and potential horizontal dissemination within the species and broader microbial communities at lower copy numbers (<2 copies per chromosome). Mobilization was inferred in all large plasmids, primarily via MOBQa-type relaxases, with differences in both replicon architecture (at least 4 different replicon types) and cargo load, which accounted for 25% to 50% of their total sequence length. Absence of the gene operons encoding the conjugative bridge in these plasmids hints on their dependency on chromosomally encoded or co-resident ICE-associated systems [15, 51, 62] for complete mobilization. In larger plasmids, stabilization systems - mostly type II TA systems, scaled with plasmid size, with approximately one TA system per 10 kb, indicating a role in maintenance under low-copy-number conditions. Collectively, insights gained on the natural architecture of native ‘*F. caldus’* plasmids reveal rules for compatibility and maintenance and offer practical guidance for synthetic vector design. For instance, in high-load plasmids, inclusion of a ParFG partitioning system and size-adjusted TA modules is likely essential to ensure stability. In contrast, small to mid-sized vectors intended for transient use may not require such systems.

To assess the genetic background of ‘*F. caldus’* strains and its relationship with plasmid carriage, occurrence and diversity of defense system (DS) and its protein complement (defensome) were evaluated in sequenced genomes and MAGs using state-of-the-art prediction tools and comparative genomics [63,64]. This analysis identified over 50 distinct DS-types across ‘*F. caldus’* genomes and MAGs, spanning well-characterized functional categories of defense [65, 66] as well as non-canonical [67] and putatively novel systems (**Fig.4a, Fig.S3**). While many of these systems are classically linked to antiviral defense, several are also known to restrict MGEs, including plasmids [20, 23–24, 68–69]. Our results confirm the presence and diversity of these systems in ‘*F. caldus’*, particularly restriction modification (RM)-systems (**Fig.4b,c**) and CRISPR-Cas systems (**Fig.4b,d**), reinforcing their potential role in shaping plasmid persistence and mobility within this species.

**Figure 4.**
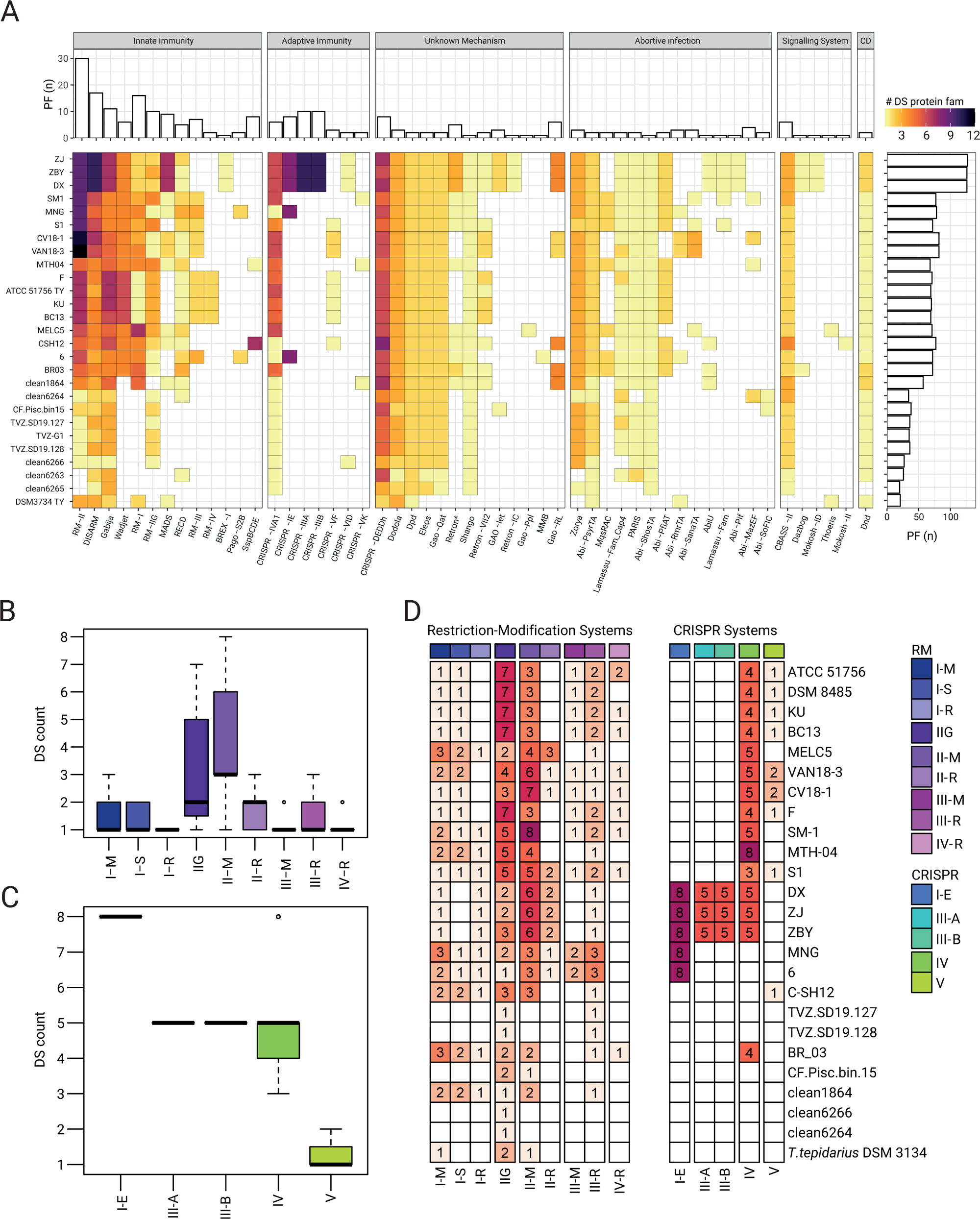
Diversity and distribution of defense systems in ‘*F. caldus’* genomes and MAGs. (**A**) Heatmap showing the presence/absence of 50 predicted defense system (DS) types across genomes and MAGs of ‘*F. caldus’*. DSs were classified and categorized after [63–66], and colored by the number of protein families (PF) identified per DS in the corresponding genomes. Core DSs (present in ≥90% of genomes/MAGs), sporadic (4–17 strains), and rare (<3 strains) distributions are highlighted. (**B-C**) Boxplot showing the number of predicted restriction-modification (R-M) systems per genome, grouped by R-M subtype (I–IV, IIG) or CRISPR-Cas system subtype (III, IV and V) across genomes and MAGs. (**D**) Heatmap displaying the distribution per strain of RM systems and CRISPR systems PFs clustered by orthology using the ProteinOrtho orthology detection tool v6.3.1 [102] at 60% identity and 60% coverage thresholds.

The abundance and distribution of DSs varied widely across strains and was generally lower in MAGs. A core set of 19 DS-types, spanning 32 genes across 16 gene neighborhoods, were present in at least 90% of genomes and MAGs (**Fig.4a**). In contrast, 21 DS subtypes were sporadically distributed across ‘*F. caldus’* strains (detected in 4 to 17 strains), while 10 were rare (present in fewer than 3 strains), with distribution patterns broadly correlating with the strains’ geographical origin or environmental source (**Table S1b**). These patterns suggest both vertical conservation and horizontal acquisition across environmental contexts. Core DSs included diverse gene clusters targeting nucleic acid degradation (e.g. Wadjet [69], BREX [70], DISARM [71]), synthesis inhibition (e.g. CBASS [72]), and phosphorothioation (e.g., Dnd [73]). Several abortive infection systems (e.g. PARIS, PsyrTA, ShosTA) and novel or composite systems (e.g. Gabija, Shango, and Zorya) were also prevalent. Of note were the type IV CRISPR-Cas and the type-I Wadjet systems implicated in anti-MGE interference, present in all strains of the species yet missing from available MAGs. In complete genomes of ‘*F. caldus’* these systems locate within, or between, know ICE [15] or within predicted ICE-like elements (data not shown), further supporting their role in the resolution of inter-MGE conflicts and/or MGE proliferation [22]. Rare or strain-specific included several location-restricted DSs such as (i) a DISARM-RM hybrid cluster (DS_7) present in the megaplasmids of SM-1 and MNTH-04, (ii) prophage-linked RM clusters (DS_8–9) associated with the AcaML1 prophage [74]; (iii) Type I-E CRISPR-Cas systems (DS_6) restricted to industrial strains from China and Africa [75]; and (iv) several integron-encoded systems (e.g., Gao-related). Less frequent DSs like BREX-I, Mokosh-II, Thoeris, among others, were detected in isolated genomes (**Fig.4**). The most frequent DS type and most abundant DS proteins were those linked to DNA restriction-modification (R-M). A total of 350 putative RM-proteins were identified across ‘*F. caldus’* genomes and MAGs, clustering into 35 distinct gene cluster arrangements. Except for two MAGs lacking RM genes, all genomes encoded 2 to 26 RM-systems, averaging 19 per genome. MAGs generally harbored fewer RM loci. No RM cluster was shared across all genomes, although the type IIG system was the most widespread, found in >85% of strains and 7 of 8 MAGs. Type II RM systems were the most abundant, comprising 19 clusters with substantial sequence diversity, occurring mostly in individual strains (**Fig.4c,d**).

Besides RM systems, CRISPR-Cas systems with known or potential relevance in plasmid interference were detected in the genomes analyzed. These included class systems of subtype IV-A, and class 2 systems of subtype V-F. Type IV was the most widespread, found in >85% of sequenced strains (and in 1 MAG), and in agreement with previously reported occurrence in the species and the *Acidithiobacillia* class [22] it is hypothesized to contribute to MGEs conflict resolution via interference between plasmids and integrative conjugative elements [76]. Interestingly, over 50% of the strains carried one or two Class 2 CRISPR-Cas type V-F (Cas12f) effectors, i.e. small (400–700 amino acids) RNA-guided endonucleases distantly related to transposon-encoded TnpB nucleases [77] that target DNA in a 5′ T-rich PAM-dependent manner, producing staggered double-stranded breaks [78], and whose collateral ssDNA cleavage capacity has been leveraged in diagnostics and gene editing [79, 80]. Beyond these roles, experimental studies with protein variants from extreme acidophiles (e.g. *Sulfoacidibacillus thermotolerans*, ex. *Acidibacillus sulfuroxidans,* AsCas12f1) have confirmed Cas12f-mediated plasmid interference in *E. coli* [78], suggesting that other orthologs of these compact effectors may also be capable of adaptive immune functions against plasmids complementing or substituting for other anti-MGE systems within the species’ broader defensome.

We next explored the global distribution patterns of *‘F. caldus’* plasmid families and defense systems across publicly available metagenomic datasets, guided by phylogenomic inference. To identify the plasmidome in local ‘*F. caldus’* populations we anchored the analysis in the plasmid backbone proteome dataset from reference and candidate plasmids obtained in this study. In the case of the defensome we used the protein repertoire recovered in sequenced strains and MAGs of the focal taxon, totalizing 243 non-redundant proteins. Among the 25 shotgun metagenomes containing representatives of the *Acidithiobacillia* class (**Table.S1b**), only 7 metagenomes exhibited ‘*F. caldus’* at relative abundances exceeding the defined filter threshold (>10%). This subset included metagenomes from geographically and environmentally diverse acidic ecosystems, including geothermal springs, volcanic river systems, acid mine drainages, and engineered biotechnological systems. Presence and relative abundance of Rep1–Rep5 replicon marker proteins across the assembled datasets (**Fig.5a**) was consistent with genome-based results generated in this study, with the pAca1.1 plasmid family being the most prevalent across samples with presence of ‘*F. caldus’* (Rep3, >75%) followed closely by the pTC-F14 family (Rep1, >60%). The metagenomic contigs recovered showed evidence of conserved backbone structures for the five plasmid families, with variations mostly confined to the plasmid cargo genes or accessory gene pool (**Table.S5**). One variant plasmid of the pTC-F14 family, displaying only 64% average similarity to the ***mob2*** module of the reference pTC-F14, was identified in the acidic riverine system of RAS-CC, expanding the repertoire of P-type mobilizable plasmids in the taxon by contributing novel gene content and organizational diversity. Also, comparative analyses of reference plasmids, and plasmid-borne contigs in draft genomes and metagenomes, provided clear evidence for the origin of larger plasmids (∼65 kb, pTcM-1) described previously in strains MNG or CSH12 [48], hinting to cointegration events of pAca1.1 and pTc-F14 family plasmids (**Fig.6a**). Stability of such large plasmids in ‘*F. caldus’* strains to be compromised, as no evidence of pCSH12 was detected in the strain (provided by Dr. D.E. Rawlings) following multiple serial passages under laboratory conditions.

**Figure 5.**
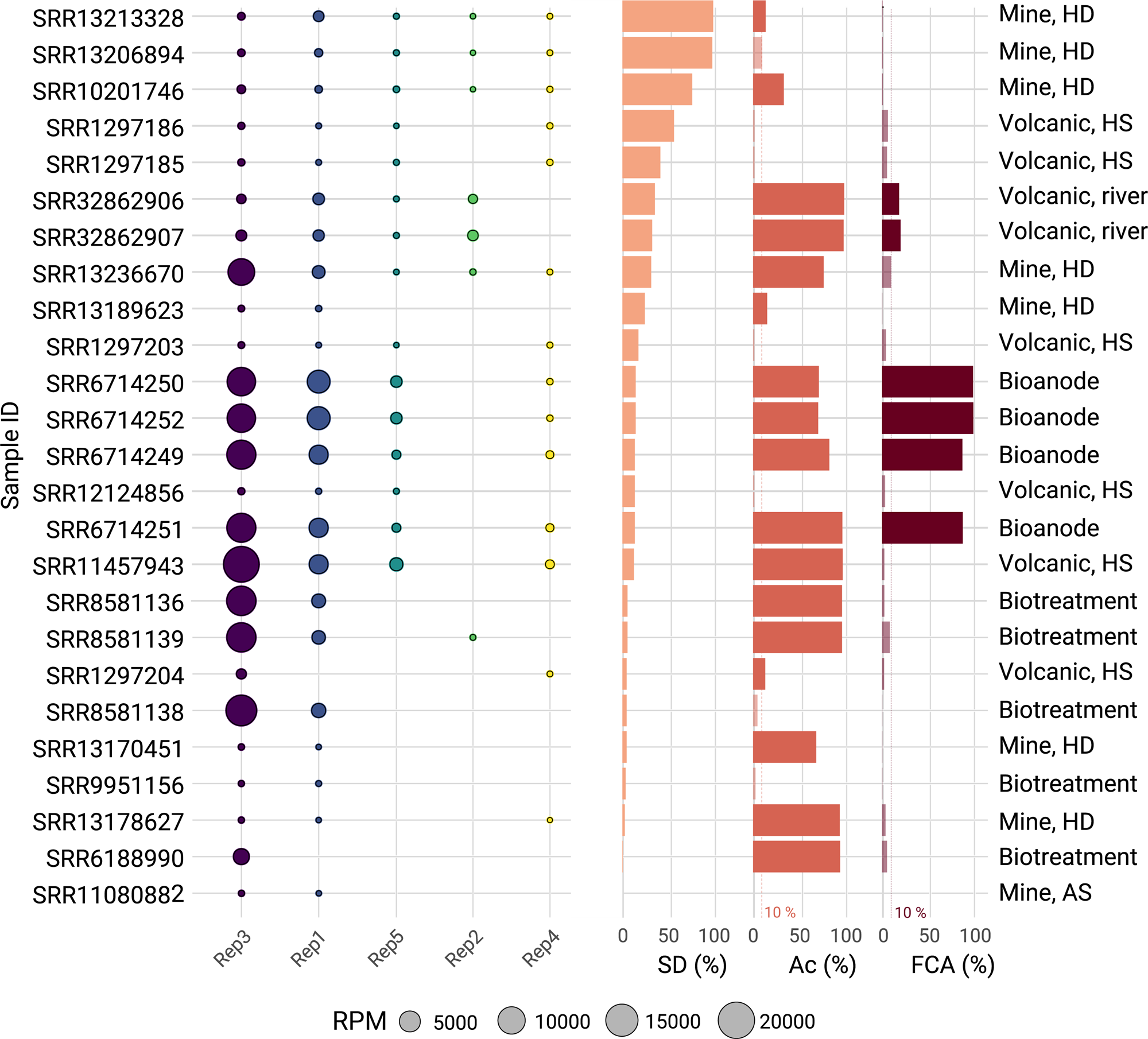
Global distribution of ‘*F. calduś* plasmid families and replicon types across acidic environments. (left) Occurrence and relative abundance (RPM) of the five ‘*F. caldus’* plasmid families (Rep1–Rep5) across 25 metagenomic samples with presence of *Acidithiobacillia* or ‘*F. caldus’* at an abundance threshold >10%, selected using the Sandpiper platform. Metagenomes were sourced from diverse acidic ecosystems, including geothermal springs, volcanic river systems, acid mine drainages, and engineered bioleaching systems (**Table S1b**). Plasmids proteins were clustered at 70% identity using CD-HIT and mapped to the metagenomes using BLAST. Replicon type was inferred by mapping against the curated plasmid backbone protein dataset derived from sequenced strains and MAGs. The abundances of *Acidithiobacillus*, ‘*F. caldus’*, and REP 1–5 proteins were calculated from read counts and normalized to relative abundances (100%). The pAca1.1-like family (Rep3) and pTC-F14-like family (Rep1) were the most widespread, consistent with their prevalence in isolate genomes. (right) Percentual abundance of ‘*F. caldus’* defense systems in the retained metagenomes (DS %), along with the relative abundance of the class (Ac %) and the ‘*F. caldus’* species (FCA %). The defense systems of ‘*F. caldus’* were evaluated by performing a BLASTp of the defensome proteins (retrieved from *Acidithiobacillus* genomes) against the metagenomic assemblies, applying a 90% identity threshold to retain high-confidence matches.

**Figure 6.**
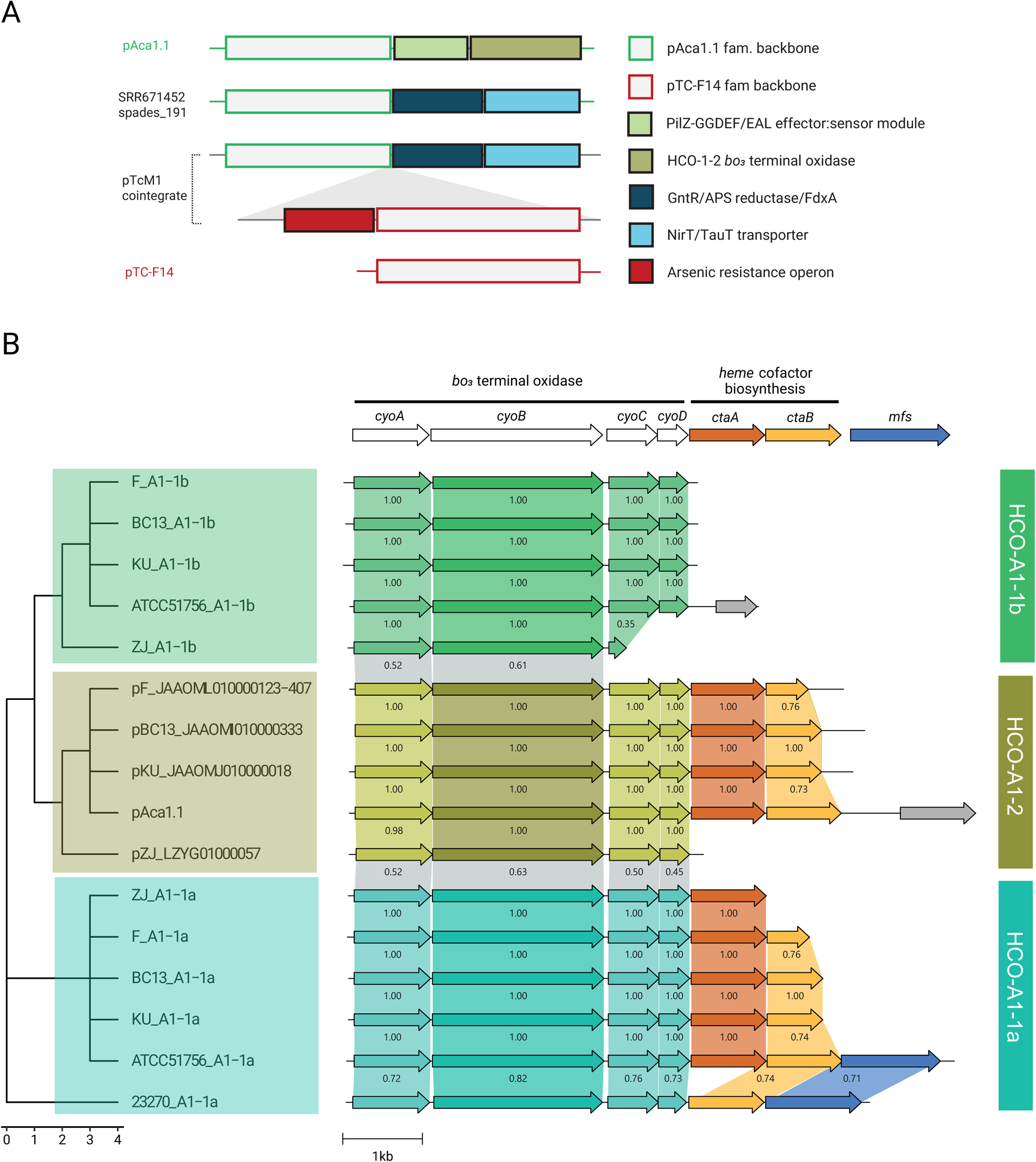
Adaptive cargo of ‘*F. caldus*’ plasmids. (A) Schematic representation showing the proposed trajectories of several cargo modules along ‘*F. caldus’* plasmids. Reference plasmid pAca1.1 and pTC-F14 backbone regions as well as adaptative cargo modules PilZ-GGDEF/EAL, HCO-A1-2 cytochrome *bo_3_* terminal oxidase, ferredoxin/APS reductase/GntR-family transcriptional regulator, ABC-type NirT/TauT sulfonate transporter and arsenic resistance operon along plasmids pAca1.1, pTcM1 and pTC-F14, and the SRR6714252_spades_191 metagenome recovered plasmid-like contig are shown. (B) Comparative analysis of HCO-A1 type cytochrome *bo_3_* terminal oxidase loci in ‘*F. caldus’* plasmids and their corresponding host strains. The cytochrome *bo_3_* terminal oxidase and linked heme cofactor biosynthesis gene neighborhoods from A1-1a And A1-1b chromosomal subunit II *cyoB* variants, along with the plasmidial A1-2 *cyoB* variants are mapped to a CyoB protein sequence-based tree, which was constructed using FastTree and visualized by using the ggtree R package. Genetic context visualizations were constructed using Clinker.js tool [115]. Names and color keys for CyoB variants are according to [14].

Plasmids from different families followed independent occurrence and abundance patterns, with all plasmid families excepting pVAN18-3 (Rep2) correlating positively with the total abundance of the host in the sample (R>0.66) and negatively with the total abundance of ‘*F. caldus’* defense systems in the sample (R<-0.35). These results suggest that, for most plasmids, the complexity of the host defensome dictates their presence and/or persistence within the host population, ultimately shaping the overall complexity of the plasmidome. In turn, for pVAN18-3 a rather complex interplay between plasmidome, defensome and environmental pressures seem to be at play. To gain further insight into these aspects, we analyzed the cargo gene complement from both genome and metagenome derived plasmid contigs. This set entailed a total of 248 non redundant proteins, 59 of which had robust functional assignments (**Table.S6a**). An additional 38 predicted proteins could be assigned to transposases (ISL3, IS4, IS5, IS30, IS66, ISChy9), and 151 protein families remained as hypotheticals. Twelve protein families with predicted function were found in both genomes and metagenomes from industrial habitats, linked to plasmid families pAca1.1, pTC-F14, pLAtc2 and pAca1.2 in decreasing frequency. In turn, 15 PFs linked to signaling, regulation and transport of nutrients, metals, and metalloids, or to competitive interactions, were only found in metagenomes (**Table.S6b**). Among PFs (n=32) exclusively associated to genomes of the species, those encoding a variant heme copper oxygen oxidase of the A1 subtype-2 (HCO-A1-2) described previously in the species [14], were the most frequent (**Fig.6b**). These were exclusively encoded in pAca1.1 family plasmids, supporting a role for this plasmid in adaptation to varying redox conditions or oxygen levels and in the evolution of HCO oxidases in the class. In contrast, pAca1.1 family plasmids from natural populations—recovered from hot spring metagenomes and engineered biotechnological systems—carried distinct cargo genes, including two operons predicted to participate in the uptake and assimilation of organosulfonate compounds as alternative sulfur sources [81]. Although sulfate is typically abundant in acidic environments, the ability of acidophiles to metabolize organosulfonates may indicate niche-specific adaptations rather than a response to sulfate scarcity. In marine ecosystems, sulfonates serve as key intermediates in trophic exchanges between phytoplankton and heterotrophic bacteria [82], raising the possibility that analogous mechanisms may facilitate community-level nutrient exchange in acidic environments. Evident presence of pAca1.1 (and pTC-F14) family plasmids in samples lacking detectable ‘*F. caldus’* (e.g. sludge metagenome, **Fig5**) further suggests that these elements may have dispersed to other members of the *Acidithiobacillia* class, reinforcing the ecological relevance of plasmid-mediated horizontal gene transfer in shaping microbial interactions and adaptability in acidic environments.

Other top-ranking COG functional categories among cargo proteins included regulation and signal transduction (**Fig.S4**). The presence of multiple GGDEF/EAL domain proteins harboring both diguanylate cyclase (GGDEF) and phosphodiesterase (EAL) domains essential for the synthesis and degradation of the second messenger c-di-GMP, alongside PilZ-type effectors [83], and key transcriptional regulators such as the flagellar master regulator FlhC [84], the multidrug efflux regulator MarR [85], and the arsenic resistance regulator ArsR [86], points to sophisticated plasmid-encoded signaling networks involved in motility, biofilm formation, and metal/metalloid resistance. Both types of traits have been linked previously to plasmids in bacteria (e.g. [87]) and in ‘*F. caldus’* [45, 51]. The documented presence of plasmid-borne arsenic resistance cassettes in ‘*F. caldus’* isolates from arsenopyrite-rich bio-oxidation plants [88], coupled with their exclusive detection in plasmids from strains inhabiting industrial environments in our dataset, highlights the niche-adaptive nature of this cargo in high-arsenic habitats and suggests host–plasmid coevolution driven by localized selective pressures.

Plasmids of the pVAN18-3 family were found in high abundance in environments where ‘*F. caldus’* was scarce (**Fig. 5**). These plasmids carried genes involved in cyclic-di-GMP signaling, a central regulatory system that controls critical adaptive traits in acidophilic bacteria, including biofilm formation, motility, cell envelope remodeling, and responses to environmental stressors [51]. In acidic, resource-limited, or fluctuating conditions, such traits are essential for microbial survival and competitiveness. The observed decoupling of plasmid abundance from that of their canonical host may arise from horizontal gene transfer to alternative hosts, environmental persistence as extracellular DNA, or stabilization mechanisms that affect the plasmid-to-host ratio, such as the host entering a dormant state. Among these scenarios, we favor the latter, considering the habitat type (riverine water column) and the environmental conditions at the sampling site (RAS-CC; pH 2.5, 18 °C), which are suboptimal for ‘*F. caldus’* and may promote dormancy or low metabolic activity in the host population. Yet, this remains to be experimentally tested. Altogether, these results suggests that pVAN18-3 plasmids, via their cyclic-di-GMP signaling cargo, may facilitate population- and/or community-level adaptation under suboptimal or fluctuating environmental conditions, and emphasize the role of plasmids as key agents in microbial resilience, horizontal gene flow, and niche expansion within extreme environments.

## Conclusions

This study provides a comprehensive analysis of the architecture and diversity of the plasmidome of ‘*F. caldus’*, an extremophilic sulfur-oxidizing bacterium inhabiting highly acidic, metal-rich and moderately hot environments. By integrating genomic and metagenomic datasets from 17 strains and multiple natural and engineered acidic habitats, we identified over 30 native plasmids belonging to 5 distinct families, defined by their unique replicon and mobilization modules. Plasmid families varied in their backbone architecture, including replication, partitioning, and stabilization systems, reflecting selective pressures favoring plasmid maintenance across populations and environmental conditions. Compatibility patterns, co-occurrence profiles, and a documented case of cointegrate formation, provided evidence of plasmid-plasmid interactions and evolutionary dynamics occurring within the species. While core features were consistent, plasmid cargo genes varied markedly between habitat types. Strains from industrial environments shared similar adaptive genes, including arsenic resistance operons and organosulfonate assimilation pathways. In contrast, environmental sequences from geothermal and volcanic sites harbored differentiated cargo, entailing different signal transduction, regulatory and transport mechanisms, relevant in the responses to redox stress or nutrient limitation. These differences point to localized host–plasmid coevolution driven by specific environmental pressures.

Although a rich diversity of plasmid-targeting defense systems was detected across genomes and metagenomes, correlation between defensome complexity and plasmid carriage was plasmid-dependent, supporting the view that adaptive cargo, not merely backbone architecture or host compatibility, plays a central role in plasmid success across ‘*F. caldus’* populations. Taken together, insights obtained in this study into plasmid compatibility, persistence, and cargo-mediated adaptation offer a conceptual and practical framework for plasmid engineering, with implications for synthetic biology, bioleaching, and bioremediation applications in extreme environments, and positions ‘*F. caldus’* as a valuable model for exploring host– plasmid–defensome interactions in extremophilic microbiomes.

## Materials and methods

### Strain isolation and growth conditions

‘*F. caldus’* isolates were obtained from acidic hydrothermal and riverine samples collected in February 2018 from multiple sites along the Copahue-Caviahue Volcanic Complex (CVCC) system, located in the Southern Volcanic Region of the Andes. Isolation was performed via direct plating and liquid culture enrichment. For direct isolation, 100 µl of water samples were spread onto solid Mineral Salt Medium (MSM) with trace elements [90] supplemented with either elemental sulfur (5 g/L) or tetrathionate (5 mM K_2_O_6_S_4_) as energy sources, and incubated aerobically at 30 °C for 15 days. For enrichment, 10 ml of water samples was inoculated into 100 ml of MSM liquid medium with matching energy sources and trace elements, adjusted to the pH 2.5 and room temperature, and incubated at 150 rpm. Once visible turbidity developed, cultures were serially diluted and plated under the same conditions. Colonies with distinct morphologies were selected and purified by repeated streaking on solid MSM. Purified strains were routinely maintained in liquid MSM medium with trace elements and a suitable energy source, incubated aerobically at 30 °C and 150 rpm, and transferred every four weeks. For biomass production cells were grown following recommended optima for ‘*Fervidacidithiobacillus caldus’* (ex. *Acidithiobacillus caldus*) [11]. Stationary phase cultures used for nucleic acid purification were processed as in [14].

### Sample collection and field procedures

Sampling was conducted at Cascada de la Culebra (CC), an acidic waterfall located in the Rio Agrio Superior (RAS) at 1,690 m.a.s.l., within the CCVC early in March 2023. Water samples were collected from the midpoint of the water column in the waterfall plunge-pool, at low-flow area of RAS-CC as in [91] (**Table.S1**). Water was pre-filtered through 8 µm Whatman grade 2 cellulose filters discs (particle-associated community) and collected on 0.22 µm MCE membrane disc filters (Millipore) for bulk water sample (25 L) metagenomic sequencing using a 500 mL Nalgene serial vacuum filtration system. Filters were stored at -20°C in the field and thereafter at -80°C until DNA extraction and sequencing.

### Genomes, MAGs and metagenomes

Genome and metagenome sequencing, as well as assembly, have been described previously [14, 92]. Public genomes and MAGs of ‘*F. caldus’* and reference strains (*Thermithiobacillus tepidarius* DSM 3134T), were obtained from the public WGS NCBI Genome database on October 2024. When available sequence read archives (SRA) for the genomes were also downloaded. For taxonomically targeted recovery of public metagenomes containing *Acidithiobacillia* class members and ‘*F. caldus’* representatives we used Sandpiper v0.3.0 [93] and downloaded the cognate files from NCBI via the SRA Run accession numbers. Quantification of *Acidithiobacillia* and ‘*F. caldus’* in each environmental sample was done via phylogenomic inference as in [92] using the BAC120 housekeeping genes matrix [94], applying identity thresholds of 85% for class-level and 95% for species-level detection, with stringent criteria of e-value < 1e-5 and query coverage 90%.

Metagenomes contributed by this study was deposited at the National Center for Biotechnology Information (NCBI) under the BioProject accession ID PRJNA914835. The complete list of genomes and metagenomes used in this study, their corresponding statistics, along with relevant metadata can be found in **Table S1**.

### Plasmid contigs identification

Downstream analyses including ORF prediction and annotation, were performed using the built-in software of the SqueezeMeta v1.6.3 pipeline [95–96] and databases GenBank, eggNOG, KEGG, and Pfam [97–100], updated in October 2023. Genomic contigs were also re-annotated using the RAST pipeline (Rapid Annotation using Subsystem Technology) [101]. Protein family (PF) clusters were assessed using the ProteinOrtho orthology detection tool v6.3.1 [102] using bidirectional BLASTp best hit and 60 identity and 60% coverage thresholds, all else as described in [14]. Similarity searches were carried out by using BLAST an PSI-BLAST algorithms [103], alignment and coverage analyses were made using Bowtie v1.2.2, and samtools v1.1 with default parameters as in [92]. To identify plasmid signatures, we assessed the presence of plasmid-signature proteins in selected contigs conducting searches against the NCBI PLSDB plasmid database [104] and by using multiple sequence alignment analysis. Target protein sequence assignments were validated against the CDD database v.3.16 using CDsearch [105], RPS-BLAST v2.2.26 and hhsearch [106], using default parameter values. Mob relaxases and other conjugative element signatures were identified and classified by using MOBscan, CONJscan and MOBFinder assignment [107 109].

### Anti-MGE defense systems prediction

General prediction and classification of defense system types in ‘*F. caldus’* genomes, MAGs, and selected metagenomes was done using DefenseFinder 11 v1.0.8 [63] and PADLOC v1.0.0 [64], applying default parameters. Individual predictions were merged via protein ID, and congruence was checked manually. Further refinement of the predictions was done using dedicated tools. CRISPR/Cas systems were reassessed by using CRISPRFinder [110], and Cas-associated proteins were classified according to established frameworks [22, 111–112]. Restriction-Modification (R-M) systems were validated through sequence similarity, conserved domains identification, and domain architecture analyses against REBASE [113]. Type-II R-M systems were also predicted by using the rmsFinder pipeline [114]. Type-II methyltransferases target sequences in plasmids and cognate chromosomes of the taxon were predicted by similarity searches using the target recognition domain (TRD) sourced from REBASE 409 (accessed in October 2024). All defense-related proteins were clustered by genomic vicinity as in [22], and manually curated to refine the defense system assignments.

### General data analysis and data visualization

Data analysis and visualization were conducted using the R language (R version 4.4.3) with ggplot2 v3.5.1, tidyverse v2.0.0, patchwork v1.3.0 and viridis v0.6.5 packages. Phylogenetic trees were visualized using the ggtree v3.12.0 R package. Genetic context and clustering visualizations were conducted using Clinker v0.0.20 program [115]

## Data availability

The biosamples used in this study are available in NCBI, under BioProject number PRJNA914835. Raw sequencing data for two metagenomes contributed in this work are available in SRA under accession numbers SRR32862906 and SRR32862907. Sample metadata and associated accession numbers are summarized in **Table S1**.

## Supplemental Material

**Figure S1. Compatibility relationships among ‘*F. caldus’* plasmids based on replicon co-occurrence. (A)** Summary matrix displaying the co-occurrences of plasmid replicon types across ‘*F. caldus’* strains. The number of plasmids per strain is indicated. **(B)** Compatibility relationships among plasmids inferred from patterns of co-occurrence across strains. All replicon types were self-incompatible, except for Rep4-type plasmids, showing up to three variant per strain. Replicon pairs Rep1:Rep3, Rep1:Rep5, and Rep3:Rep4 frequently coexisted, with the latter being the most prevalent combination. Rep3:Rep5 concurrence was infrequent and Rep2 with any other replicon were mutually exclusive, suggesting functional incompatibility. One cointegrate plasmid carrying two replicons (Rep1 and Rep3) was also identified, suggesting recombination between plasmids of different families.

**Figure S2. Predicted origin of transfer (*oriT*) of mobilizable plasmids in ‘*F. calduś*.** Predicted *ori*T regions were consistently located between the *mobA* relaxase gene and the divergently transcribed RAPs, regardless of the *mob* module. Sequence conservation was high among plasmids of the same mobilization module type but variable between types. All oriTs conformed to a palindromic structure adjacent to a predicted nick site, resembling *ori*Ts from plasmids pTF1 and pTC-F14 [17, 54]. In bold conserved bases between plasmids of the same mob type and in red the inferred nick site.

**Figure S3. Defense systems diversity and distribution in ‘*F. caldus*’ genomes.** (**A**) Frequency histogram showing the number of defense systems (DS) that could be assigned to a single genomic locus, and therefore clustered as “defense islands”. (**B**) Number of DS-associated unique protein families (PFs), and their prevalence in ‘*F. caldus’* genomes. The prevalence was defined as the number of genomes in which a given DS PF was identified divided by the total genomes analyzed. (**C**) Total number of DS-associated PFs identified in each ‘*F. caldus’* genome sequence, ranked from highest to lowest. PFs were defined by clustering orthologs using ProteinOrtho [102] with thresholds of 60% identity and 60% coverage. Defense systems were classified after ProteinOrtho [63–67].

**Figure S4. Functional assignment of ‘*F. caldus’* plasmid encoded protein cargo**. Protein sequences recovered from non-backbone plasmid regions were used as queries for homology searches using the COG database by using eggNOG. Number of coding sequences assigned to a given COG top level categories.

## Supplementary Tables

**Table S1. Genomes and metagenomes used in the analysis of the plasmidome and defensome of ‘*F. caldus’*.** (**A**) Sequenced strains and MAGs used in this study. (**B**) List of metagenomes recovered from the SRA database (NCBI). Only metagenomes passing the inclusion threshold (presence of *Acidithiobacillia* and/or >10% ‘*F. caldus’*) were analyzed. Basic meta/genome statistics and descriptive metadata were retrieved from NCBI and/or listed literature.

**Table S2. Key attributes of reference and candidate plasmids in ‘*F. caldus’* genomes.** (**A**) Sequence-derived and functional data summary for plasmids and plasmid-contigs affiliated with the five plasmid families (PLASMI_fam) identified in ‘*F. caldus’* strains and metagenome-assembled genomes (MAGs) to support comparative analyses of plasmid architecture, diversity, copy number, and compatibility. Each entry includes the plasmid name, host strain or MAG, and unique plasmid or contig identifier. The replication (REP) and mobilization (MOB) modules are annotated, along with the presence of partitioning (PAR) and toxin-antitoxin (TA) systems, including the number of TA modules and predicted target type (e.g., mRNA, tRNA, EF-Tu). Plasmid status is indicated as full or assembled based on assembly continuity. Genomic features include the plasmid size (in base pairs), the average G+C content of plasmid contigs, and the chromosomal G+C average for the respective host genome. The absolute difference in G+C content (Δ G+C, %) between plasmid and host chromosome is shown as an indicator of potential foreign origin. Coverage statistics include average depth of coverage for plasmid and chromosomal contigs, and calculated relative plasmid copy number (fold-abundance). Functional annotation metrics include total predicted genes and coding sequences (CDSs), and the relative proportion of core (shared) vs. cargo (variable/adaptive) genes, based on protein clustering. (**B**) Sequence-derived and functional data summary for the cognate chromosomes in ‘*F. caldus’*strains and metagenome-assembled genomes (MAGs).

**Table S2. Key attributes of reference and candidate plasmids in ‘*F. caldus*’ genomes.** (**A**) Sequence-derived and functional data summary for plasmids and plasmid-contigs affiliated with the five plasmid families (PLASMID_fam) identified in ‘*F. caldus’* strains and metagenome-assembled genomes (MAGs) to support comparative analyses of plasmid architecture, diversity, copy number, and compatibility. Each entry includes the unique plasmid or contig identifiers and basic genomic statistics. Annotated replication (REP) and mobilization (MOB) modules, along with the presence of partitioning (PAR) and toxin-antitoxin (TA) systems, including the number of TA modules is indicated. Plasmid status is indicated as full or assembled based on assembly continuity. Genomic features include the plasmid size (in base pairs), the average G+C content of plasmid contigs, and the chromosomal G+C average for the respective host genome. The absolute difference in G+C content (Δ G+C, %) between plasmid and host chromosome is shown as an indicator of potential foreign origin. Coverage statistics include average depth of coverage for plasmid and chromosomal contigs, and calculated relative plasmid copy number (fold-abundance). Functional annotation metrics include total predicted genes and coding sequences (CDSs), and the relative proportion of core (shared) vs. cargo (variable/adaptive) genes, based on protein clustering. (**B**) Sequence-derived and functional data summary for the cognate chromosomes in ‘*F. caldus’* strains and metagenome-assembled genomes (MAGs).

**Table S3. Classification, gene content, and species-level distribution of plasmid functional modules in ‘*F. calduś* strains and MAGs.** Backbone-based plasmid family classification was supported by functionally coherent **replication and mobilization modules.** Each entry represents a unique plasmid-borne protein family, classified by plasmid family (PLASMID_TYPE), module type (MODULE_TYPE), with further subtype designation (MODULE_SUBTYPE). For each backbone protein family, the corresponding cluster names (CLUSTER_NAME) are listed. Columns (corresponding to genome assemblies) indicate the presence or absence of each PF in individual strains or MAGs, by the protein_ID.

**Table S4. Genome-derived plasmid contigs and family assignments.** Summary of plasmid contigs identified in ‘*F. caldus’* genomic datasets analyzed, including contig IDs, sample origin, plasmid family (based on Rep-type), backbone organization, and cargo proteins.

**Table S5. Metagenome-derived plasmid contigs and family assignments.** Summary of plasmid contigs identified in metagenomic datasets analyzed with presence of ‘*F. caldus’*, including contig IDs, sample origin, plasmid family (based on Rep-type), backbone organization, and cargo proteins. Sample metadata is listed in **Table S1b**.

**Table S6. Defense systems identified across ‘*F. caldus’* genomes and MAGs.** Presence, classification, and frequency of predicted defense systems (DSs) across genomes and metagenome-assembled genomes (MAGs) of the species. Each row corresponds to a distinct defense system cluster (DEFENSOME), categorized into known system types (DS_system_d) such as restriction-modification (R-M), CRISPR-Cas, DISARM, BREX, and others, following the classification frameworks established in [63–66]. Systems were grouped by prevalence into core (present in ≥90% of genomes/MAGs), sporadic (found in 4–17 strains), or rare (in <3 strains). Higher-level functional grouping of each system (e.g., DNA targeting, abortive infection, composite). For each system, the gene name is included, along with the most common defense system name, and annotated function, representing dominant hits in orthologous clusters. Protein families associated with each DS, was clustered using ProteinOrtho [102], with a 60% identity and 60% coverage threshold.

**Table S7. Functional annotation of plasmid cargo genes.** List of all non-redundant protein families with assigned function (n=59) identified in the ‘*F. caldus’* plasmid cargo across genomes and metagenomes including sequence identity based predicted function, COG/KEGG/PFAM assignments and occurrence in the analyzed datasets

## Acknowledgements

This work was supported the Agencia Nacional de Investigación y Desarrollo (ANID) under Grants ANID/FONDECYT 1221035 (RQ), 3230527 (FI), 1230217 (BD), ANID/Exploración 13220230 (SB, RQ), ANID/BASAL/FB210008 (RQ); ANID/Millenium Institute CN2021_044 (BD), ANID/PhD Scholarships (CR-V 10218491, AZ-A 21242020), Competition for Research Regular Projects, year 2023, code LPR23-09, Universidad Tecnológica Metropolitana, and Vicerrectoría de Investigación y Doctorados - Universidad San Sebastián, PhD scholarships 10202936 (SP-A), 80015086 (GC-T), Postdoctorate scholarship USS-FIN-23-PDOC-03 (AM-B) and field trip travel grants USS-FIN-23-PASD-04 (SP-A). The funders had no role in study design, data collection and interpretation, or the decision to submit the work for publication. The authors do not have any conflict of interest. The authors thank the authorities of the Provincial Thermal Baths Agency (EPROTEN) and the Directorate of Protected Natural Areas (ANP) of the province of Neuquén Argentina for allowing access and sampling in the Copahue-Caviahue Provincial Park and Rubén Vargas (alias Caniche) for guidance during ascent to the Copahue volcano. Yasna Gallardo and Hector Carrasco aided with sampling, microbiological and molecular biology analyses.

